# Environmental Drivers of Calling Activity in a Southern Subtropical Anuran Assemblage: Insights from Passive Acoustic Monitoring

**DOI:** 10.1101/2025.08.26.672434

**Authors:** Paula Pouso, Álvaro Cabana, Clara Nieto Methol

## Abstract

In anurans, acoustic communication is a crucial reproductive behavior shaped by individual traits, social interactions, and environmental cues. External factors such as temperature, rainfall, and photoperiod can elicit physiological responses that drive behavioral rhythms at individual and population scales. While temperature and rainfall effects on anuran calling activity are well-established, photoperiodic influences remain comparatively understudied, particularly within southern subtropical assemblages.

We assessed hourly calling activity of *Boana pulchella* in Uruguay over a 12-month period using Passive Acoustic Monitoring (PAM) and validated semi-automated data processing methods to examine environmental drivers in a subtropical permanent pond. Calling behavior increased during the warmer months, showing a clear seasonal pattern. Peak activity timing remained consistent across seasons, but the temporal window of vocalizations expanded during winter (long nights) and contracted during summer (long days), reflecting photoperiodic variation.

Linear regression analyses showed significant effects of photoperiod, temperature and their interaction on calling activity, while rainfall and atmospheric pressure showed no effect. These findings underscore the regulatory role of photoperiod in shaping reproductive acoustic behavior and highlight the need to further explore its physiological and adaptive significance, especially within underrepresented subtropical assemblages.

## Introduction

Physical factors such as temperature, rainfall, and photoperiod trigger physiological responses that drive behavior at both individual and population levels, shaping key life-history events (Chmura et al., 2019; Visser et al., 2010). Vertebrate assemblages have been extensively studied for their adaptability in vocal behavior in response to environmental conditions, particularly in reproductive contexts (Gerhardt, 2001; Hawkins et al., 2025; Oestreich et al., 2024), for instance, in marine mammals (Åsvestad et al., 2024), birds, (Hao et al., 2025), lizards (Lin et al., 2024; Young et al., 2013), and frogs (Brodie et al., 2025).

The drivers of acoustic signal emission—ranging from behavioral dynamics to neurophysiology—have been extensively studied across a wide range of vertebrate taxa (Simmons and Fay, 2006). However, most research has focused on individual-level processes (Barkan et al., 2021; Brenowitz et al., 1985; Gentner and Margoliash, 2003). Chorusing anuran species provide an excellent model for studying acoustic behavior at the population level and its regulation by environmental cues (Narins et al., 2023; Navas, 1996). It is well established that physical factors play a significant role in regulating chorusing activity (Duellman and Trueb, 1994). For example, the number of calling individuals correlated positively with temperature in a Colombian dendrobatid, *Colostethus subpunctatus* (Navas, 1996). Calling activity was strongly associated with rainfall in an Australian microhylid, *Austrochaperina robusta* (Hauselberger and Alford, 2005) and humidity in a Canadian ranid, *Rana clamitans* (Feng et al., 2009; Narins and Meenderink, 2014a; Oseen and Wassersug, 2002; Pauly et al., 2006). However, the influence of photoperiod on calling activity in chorusing anurans has been less studied (Both et al., 2008; Canavero and Arim, 2009; Schalk and Saenz, 2016). Moreover, for many decades, the study of environmental cues affecting acoustic signal emission has focused mainly on tropical and temperate anuran populations, leaving southern subtropical populations understudied (Both et al., 2008; Canavero et al., 2008).

Historically, acoustic recordings of anurans has relied on active monitoring methods (Feng et al., 2009; Narins and Meenderink, 2014b; Oseen and Wassersug, 2002; Pauly et al., 2006), and much of this research assessed calling activity in a fragmented fashion, often restricted to specific times of day or seasons (Canavero et al., 2008; Canavero and Arim, 2009). The advent of passive acoustic monitoring (PAM) devices has recently enabled continuous data collection over extended periods (Gibb et al., 2019; Ribeiro et al., 2017; Sugai et al., 2021). This technological advancement facilitates a more detailed analysis of acoustic signaling phenomena and their correlation with other simultaneously recorded variables. It allows for the development of increasingly accessible and precise approaches to characterize the complex acoustic environments where communication occurs. Nonetheless, despite the benefits of PAM, automated processing remains challenging because of the substantial volume of data collected. Although vocal activity in tropical frogs has been studied using PAM, chorusing and its environmental regulation in subtropical species in South America, has seldom been assessed using PAM (Bonnefond et al., 2020; Boullhesen et al., 2019; Duarte et al., 2019).

*Boana pulchella* is a widely distributed hylid species in southern South America, exhibiting a prolonged reproductive strategy typical of temperate amphibians (Maneyro and Carreira, 2012). Males form choruses to attract females and facilitate reproductive activities. *B. pulchella* males vocalize with three distinct note types that vary in frequency and duration: the “doublet”, formed by notes 1 and 2, functions as the primary and most frequent advertisement call (see Figure 1D), while the less common “squeak,” composed of note 3, occurs less frequently (Basso and Basso, 1987; Rodriguez-Santiago et al., 2023). Although the species has a relatively simple vocal repertoire, it exhibits notable flexibility and variability in its vocal emission patterns, which are influenced by social context (Rodriguez-Santiago et al., 2023). While some studies have investigated the calling activity of *B. pulchella* choruses, data collection has relied on active and sporadic monitoring, typically involving multiple species and exploring only a limited set of environmental cues (Canavero et al., 2008).

**Figure 1.**
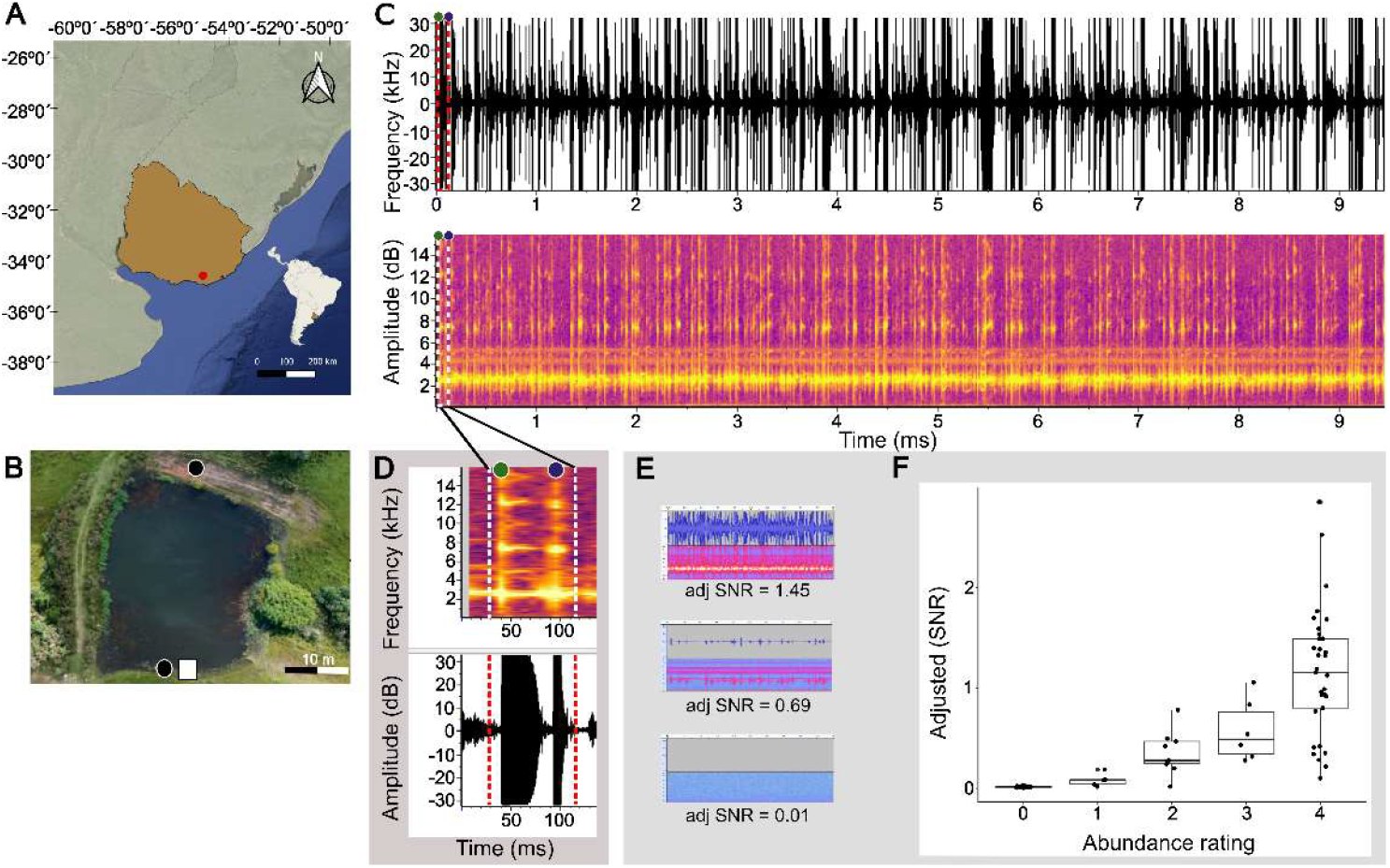
Data collection and manual validation of calling activity in a *Boana pulchella* assemblage. (A) Location of the study site in Maldonado, Uruguay (red dot), where passive acoustic monitoring (PAM) was conducted. (B) Aerial view of the permanent pond. Black circles indicate the positions of PAM dataloggers used to record calling activity; the square denotes the location of environmental sensors for light intensity and temperature. (C) Representative sample recording from a PAM datalogger capturing calling activity within the assemblage. Top: oscillogram. Bottom: spectrogram. (D) Close-up view of the spectrogram shown in (C), highlighting a characteristic “doublet” advertisement call produced by a *Boana pulchella* male. Bottom: close-up view of the oscillogram shown in (C). “Note 1” is indicated by the green dot, and “Note 2” by the blue dot. (E) Examples of PAM audio recordings and their corresponding adjusted SNR values (F) Manual validation of SNR scores based on audio file selection. A positive correlation was observed between the vocal abundance rating and the adjusted SNR values (Spearman’s ρ = 0.89, p<.001).

Our study investigates the influence of temperature, rainfall, and photoperiod on the seasonal and daily calling activity of an anuran assemblage in a southern subtropical region, using passive acoustic monitoring (PAM). Our aims were to: (1) describe the seasonal and daily patterns of calling behavior within the *B. pulchella* assemblage; (2) validate the semi-automated processing of a large acoustic dataset; and (3) assess the relationships between environmental variables and calling activity in a *B. pulchella* population in its natural context.

## Materials and Methods

### Data collection

We conducted daily recordings of chorus calling activity, temperature, and light intensity at Paraje Zanja del Tigre, Maldonado, Uruguay (34°32’47.8”S, 55°05’18.8”W) from March 16 to October 28, 2020, and from October 16, 2022 to March 20, 2023, (Figure 1A). Climatic conditions in the study area correspond to a humid subtropical regime, with warm, humid summers and cool, humid winters. The mean annual temperature was 21.5 °C, ranging from 1.2 - 34.8 °C. Annual precipitation ranged from 2.5 - 54.5 mm, with a relatively uniform distribution throughout the year.

We used two types of data collection devices positioned in a permanent pond inhabited by *B. pulchella* (Figure 1B). First, autonomous audio recording units (ARU) for acoustic recordings (AudioMoth v1.1.0, Open Acoustic Devices), equipped with omnidirectional microphones were mounted on trees approximately 1.5 m above the ground and protected from rain with a resealable plastic bag. Each device was configured to capture a 10 s audio clip every 10 min throughout a 14 h daily recording window (5:30 PM - 7:30 AM), yielding 18,844 recordings in total. This setup offers fine temporal resolution suitable for long-term acoustic monitoring (Cañas et al., 2023; Maneyro and Carreira, 2012; Ziegler et al., 2011). A sample recording is shown in Figure 1C, along with a magnified view of a doublet call (Figure 1D). Recordings were made in mono at a sampling rate of 32 KHz with 32-bit resolution. It is worth noting that our study did not include daytime chorus activity. However, previous research on this species indicates that vocal activity occurs exclusively at night (Canavero et al., 2008).

Second, a temperature and light intensity data logger (HOBO UA-002-64, Onset Computer Corporation, Bourne, MA, USA) was mounted on a pole ∼1 m above the ground, continuously exposed to ambient light. Measurements were recorded hourly over a 24 h period each day. Finally, we obtained other climate data (relative humidity, atmospheric pressure and daily rainfall) from Uruguay’s national weather service (INUMET) recorded at the Laguna del Sauce International Airport weather station, located 30 km from the study area.

To simplify data analysis, we grouped data samples by night. Specifically, data recorded at 11:50 PM were grouped with data from 12:00 AM the following day. To achieve this, we shifted the date transition time from 12:00 AM to 3:00 PM. As a result, all data points from 12:00 AM to 2:59 PM were assigned the same date as those from 3:00 PM to 11:59 PM of the previous day.

All data processing and statistical analyses were performed using the R programming language (R Core Team, 2025) in RStudio (Posit team, 2025).

### Environmental data processing

We calculated the proportion of data points exceeding 100 lux (photoperiod phase), as well as the daily averages of temperature, relative humidity, and atmospheric pressure. A 7d rolling average was also applied to daily precipitation values to estimate recent rainfall accumulation. In some analyses, photoperiod phase was further categorized into tertiles based on its distribution across all days.

### Audio data processing

To automatically quantify vocal activity level of *B. pulchella* in a natural context using passive monitoring devices, we analyzed the spectral properties of the calls. Figure 1 C shows an oscillogram and a spectrogram of a typical audio clip, in which individual calls are distinguishable, and Figure 1 D shows a close-up showing two individual notes comprising a doublet. Individual notes have a fundamental frequency around 2.5 KHz, with contextual and individual variability related to environmental and physiological factors (Ziegler et al., 2016, 2011).

During periods of high vocal activity, notes from different individuals within a chorus overlap rendering note tracking unfeasible. Thus, we chose to quantify the power spectral density of the audio signal at a frequency band centered at the fundamental frequency of the doublets. Since non-specific noise in the environment could contribute energy to this band, we used as a specific indicator of vocal activity the signal to noise ratio (SNR). For each audio file, we obtained the power spectral density for a 4s interval starting after 1s using the *seewave* package (Sueur et al., 2008). We computed the SNR as a ratio between median power spectral densities for the vocalization frequency range (2 to 3 KHz), relative to a baseline noise frequency range (.3 to 1.3 KHz). Throughout the recording period, we used two devices in three different recording periods, which resulted in changes in the position and orientation of the devices. This resulted in SNR that significantly differ in value, which is especially evident when comparing data from consecutive nights that fall in different recording periods. To ensure the compatibility of these measures, we normalized the SNR by dividing its value by the 90th percentile of the corresponding recording period (which varied from 31.0 to 48.9).

### Manual validation of SNR measures

Since most audio files do not register vocal activity, we sought to obtain a balanced validation sample comprised of recordings with and without activity. Hence, we divided the whole dataset around an empirical SNR threshold value of 0.7 that roughly distinguishes periods with and without vocal activity. Then, we randomly selected 50 files from each division for manual validation. The resulting sample had a median SNR value of 0.06, with an interquartile range (IQR) of 0.84.

The validation sample was then manually inspected independently by three judges that were blind to the corresponding SNR values. For each audio file, the judges assigned a rank abundance score (see (Canavero et al., 2008)) as follows: 0 (no calling activity), 1 (one calling male), 2 (two or three calling males), 3 (more than three calling males, with individual calls clearly distinguishable), and 4 (a chorus, in which individual calls are no longer distinguishable). Abundance scores from the three judges were highly correlated (all pairwise Spearman’s coefficients ρ = 0.98, p<.001). A consensus abundance score was constructed by averaging the individual abundance scores and rounding to the nearest integer.

The resulting consensus abundance scores and SNR values were highly correlated (Spearman’s ρ = 0.89, p<.001). In recordings where no calls were identified, SNR values were below 0.04. Overall, SNR is highly indicative of the overall level of vocal activity (see Figure 1F). However, some recordings with the highest abundance score (corresponding to full-blown choral activity) show SNR values characteristic of abundance scores 2 or 3 (two or three distinguishable calling males). Close inspection suggests that an increased noise baseline amplitude leads to a reduced SNR in these samples. Despite this shortcoming, our SNR measure is a reliable indicator of vocal activity.

### Statistical analyses

To explore seasonal differences in hourly calling activity, we first restricted the analysis to nights with substantial chorus activity, excluding those in which the maximum SNR value was below 1. We then selected all audio recordings that contained choral activity, defined as having an SNR value above 0.5, and grouped them according to photoperiod tertiles. For each group, we calculated the median and interquartile range (IQR) of the timestamps of the selected recordings. To test differences between groups, we conducted randomization tests using pairwise differences in these indicators (median and IQR) as test statistics. In each iteration, photoperiod labels were randomly permuted across recordings, group medians or IQRs were recalculated, and the resulting pairwise differences were stored. After 10,000 iterations, p-values were obtained by comparing the observed pairwise differences to the distribution generated under the null hypothesis of random group assignment. A similar procedure was followed to test seasonal differences in the time of peak chorus activity.

The influence of environmental factors on calling activity throughout the year was evaluated using multiple regression in which the environmental factors were included as predictors, and the calling activity as the response variable. To reduce dependencies between data from consecutive days, we subsampled the dataset by retaining one out of every five consecutive days, resulting in a final set of 90 observations instead of 359. We evaluated and compared models using R^2^, AIC and BIC statistics.

## Results

### Calling activity varies across seasonal cycles and nocturnal hours

Calling activity predominantly occurs from mid-August to late April, corresponding to the end of winter and the end of autumn in the Southern Hemisphere (Figure 2A). Vocalization is minimal from May to late August, but it was exhibited in July, coinciding with a temporary increase in temperature during that period (Figure 2A, Supplementary Figure S1). The daily onset and termination of calling activity are coincide with sunset and sunrise respectively, spanning from 7 PM to 5 AM in winter (late August to November) and 9 PM – 3 AM in summer (Figure 2A) Peak calling activity occurs around 11 PM across seasons, showing no significant seasonal variation (Supplementary Figure S2, p-values >.4).

**Figure 2.**
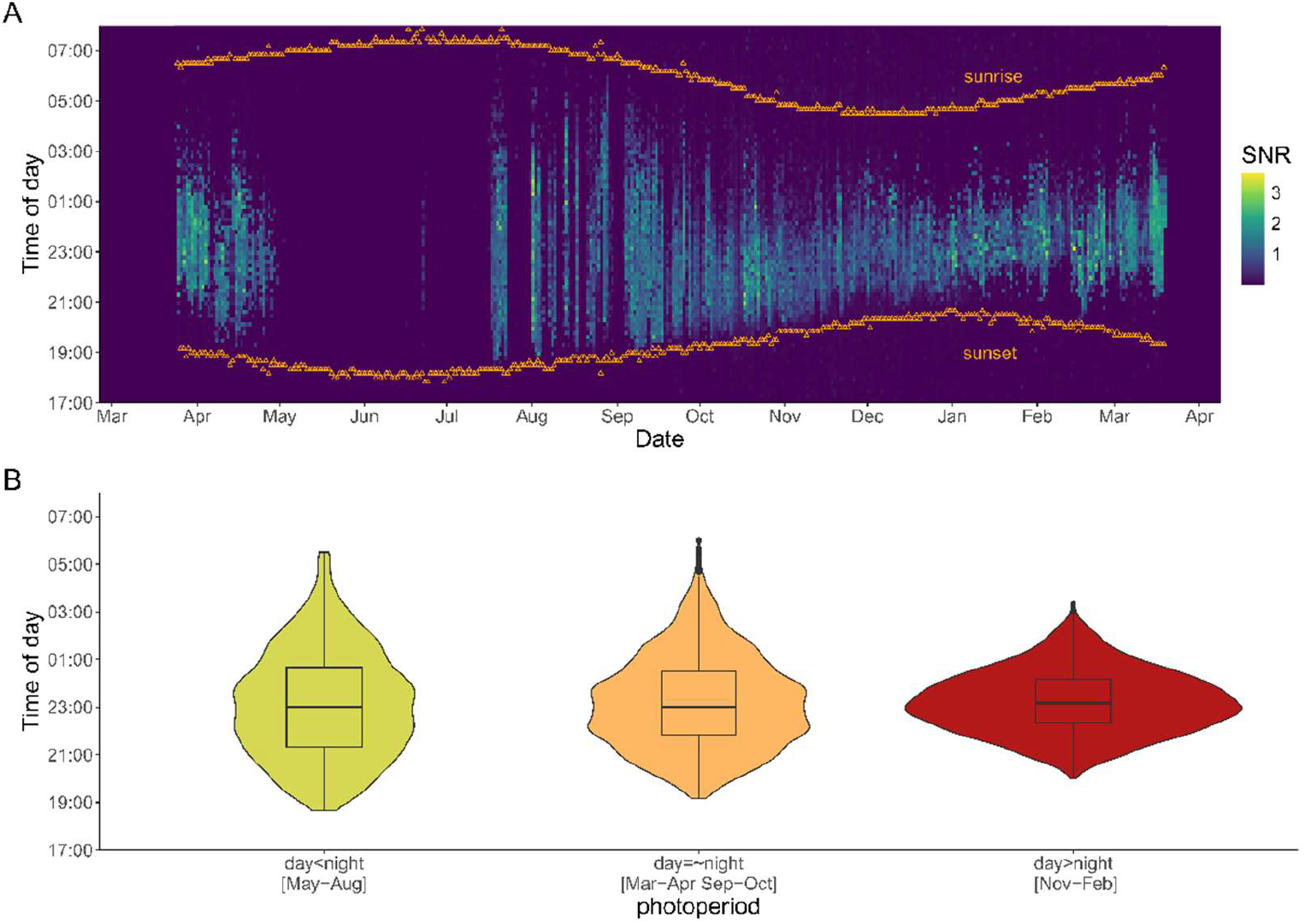
Hourly calling activity of the *Boana pulchella* assemblage throughout the year. (A) Hourly calling activity throughout the year, represented across months. Calling activity is prolonged throughout the year but ceases during certain months. Additionally, its temporal distribution varies seasonally, with vocalizations constrained to specific time windows depending on the time of year. Colors depict the signal to noise ratio (SNR). For simplicity, dates were ordered irrespective of the calendar year to which they actually belong. (You probably want to change the months on the x-axis to English or to numbers (i.e., 1= January, 2= Feb, etc.) (B) Calling activity distribution under different photoperiod conditions. The temporal distribution of calling activity exhibits seasonal variation, with a defined time window that changes throughout the year, being wider in long nights and shorter during long days (p<.001), with a constant median at around 23:00 hs (p>.6). (On the x-axis, you might include the months for each period, for example: day<night (July-Oct) day > night (Nov – Mar) etc. SNR: Signal-to-noise ratio.

The nightly duration of activity changes according to season and photoperiod (Figure 2B). During winter, when daylight is shorter than nighttime, calling extends over a broader window time (median: 11:00 PM; IQR: 200 minutes). In contrast, when daylight exceeds nighttime during summer, calling activity is confined to a narrower time window (median: 11:17 PM; IQR: 110 minutes; p<.001). Around the equinoxes, when day and night lengths are equal, the time window is broader than in winter but narrower than in summer (median: 11:00 PM; IQR: 160 minutes; p<.001).

### Differential Effects of Environmental Factors on Calling Activity

To explore the effects of environmental variables on calling activity, we first calculated Spearman correlation coefficients between all variables (Table 1). Only photoperiod was significantly correlated with calling activity (r = 0.46, p <.001), although photoperiod itself was also significantly correlated with air temperature and cumulative rainfall. Air temperature, in turn, showed a strong negative correlation with atmospheric pressure. These correlations suggest potential multicollinearity, which was considered in model selection.

**Table 1.** Spearman correlations between SNR and environmental variables. *: p < 0.05; **: p < 0.01; ***: p < 0.001. SNR: signal-to-noise ratio.

|  | Photoperiod | Air Temperature | Cumulative rainfall | Atmospheric pressure | Relative humidity |
| --- | --- | --- | --- | --- | --- |
| **Calling activity (mean SNR)** | 0.46*** | 0.27 | -0.14 | -0.12 | 0.12 |
| **Photoperiod** |  | 0.45*** | -0.29* | -0.17 | -0.23 |
| **Air Temperature** |  |  | -0.02 | -0.57*** | 0.19 |
| **Cumulative rainfall** |  |  |  | -0.08 | 0.19 |
| **Atmospheric pressure** |  |  |  |  | -0.11 |

We then tested a series of linear regression models with environmental variables as predictors and calling activity (mean SNR) as the response variable (Table 2). Model 0 included all five predictors. Based on visual inspection of the data (Figure 3B), we hypothesized an interaction between photoperiod and air temperature: during shorter nights, higher temperatures were associated with increased calling activity, whereas during longer days, higher temperatures appeared to reduce calling activity. This motivated the inclusion of an interaction term in Model 1, which resulted in the largest improvement in model fit overall. No interaction between photoperiod and cumulative rainfall was observed (Figure 3A).

**Table 2.** Model fit indicators for five regression models tested. The best-fitting model is highlighted in bold. AIC: Akaike information criterion. BIC: bayesian information criterion.

| model | terms | Adj. R<sup>2</sup> | AIC | BIC |
| --- | --- | --- | --- | --- |
| Model 0 | photo + temp + rain + atm_pressure + rel_hum | 0.19 | -9.49 | 8.01 |
| Model 1 | photo + temp + rain + atm_pressure + rel_hum + photo×temp | 0.36 | -30.77 | -13.27 |
| Model 2 | photo + photo<sup>2</sup> + temp + rain + atm_pressure + rel_hum + photo×temp | 0.47 | -46.89 | -24.39 |
| Model 3 | photo + photo<sup>2</sup> + temp + rain + photo×temp | 0.44 | -43.65 | -26.15 |
| **Model 4** | **photo + photo<sup>2</sup> + temp + rel_hum + photo×temp** | **0.47** | **-47.68** | **-30.18** |
| Model 5 | photo + photo<sup>2</sup> + temp + atm_pressure + photo×temp | 0.45 | -45.10 | -27.60 |

**Figure 3.**
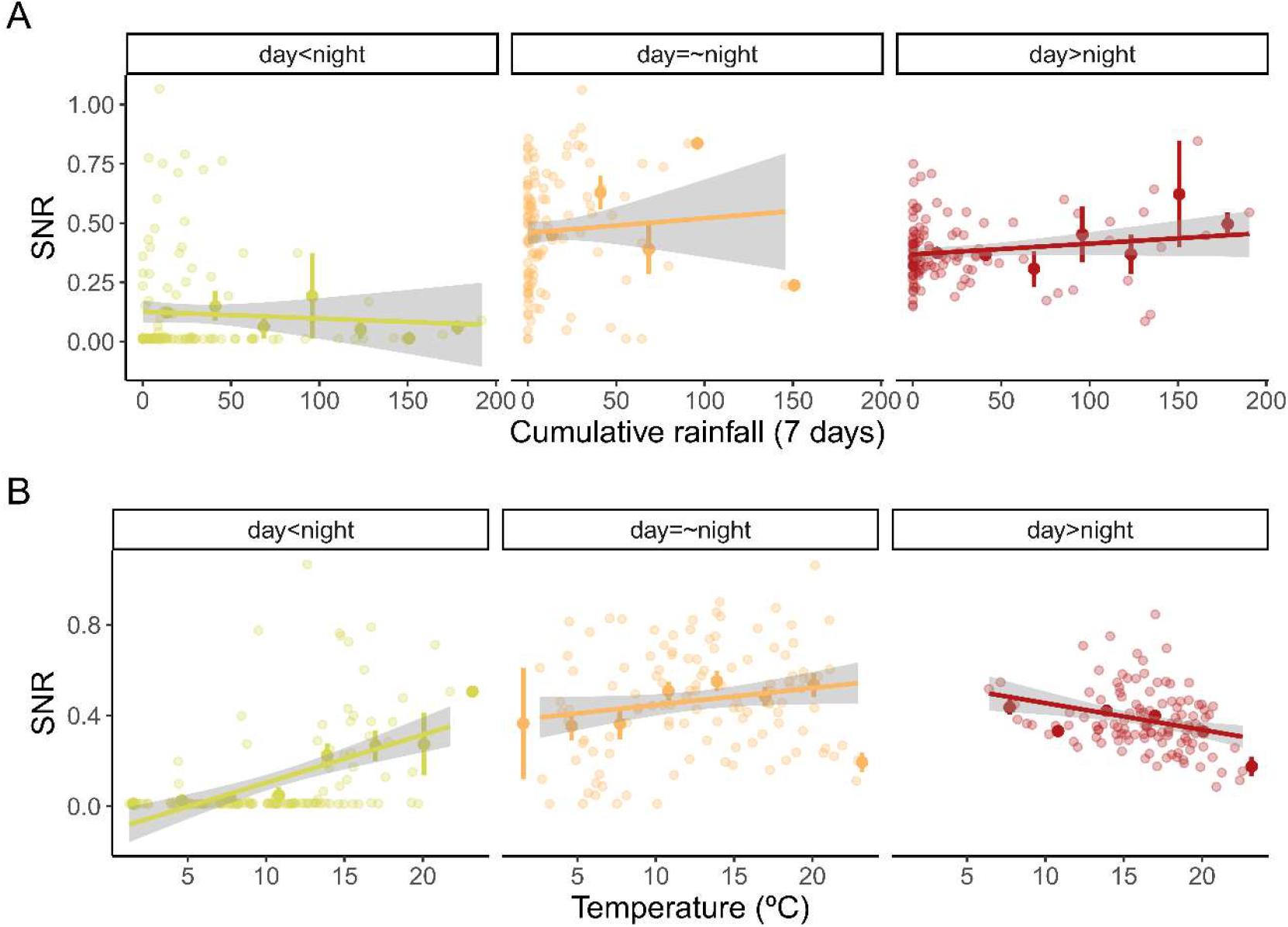
Effect of cumulative rainfall and temperature on calling activity (SNR) in relation to photoperiodic variation. (A) Relationship between SNR and cumulative rainfall across photoperiod. No significant association was found between cumulative rainfall and SNR across photoperiodic conditions (p>.05). (B) Relationship between SNR and temperature across photoperiod. A significant association was found between temperature and SNR across photoperiodic conditions (p<.05). SNR: signal to noise ratio. The photoperiod is measured by the duration of daylight and nighttime in hours.

Further inspection also suggested a nonlinear relationship between photoperiod and calling activity, prompting the addition of a quadratic term in Model 2. Since several predictors in Model 2 were either non-significant or marginally significant, we tested three simplified models (Models 3–5), each including only one of the weaker predictors: cumulative rainfall, relative humidity, or atmospheric pressure, respectively.

Among all models tested, Model 4 achieved the highest adjusted R^2^ (0.47) and the lowest AIC and BIC values, indicating the best fit. In all models, temperature, photoperiod, and their interaction were statistically significant. Cumulative rainfall was not significant in any model, and atmospheric pressure only reached significance sporadically.

Table 3 summarizes the parameter estimates for Model 4. Photoperiod and temperature had positive effects on calling activity, but the significant negative interaction term indicates that the positive effect of temperature diminishes as photoperiod increases. Additionally, the negative coefficient for the quadratic term of photoperiod implies that the increase in calling activity associated with longer days follows a decelerating trend. Finally, relative humidity showed a small but significant positive effect on calling activity (p =.04), suggesting that increased humidity may slightly enhance vocal behavior.

**Table 3.**
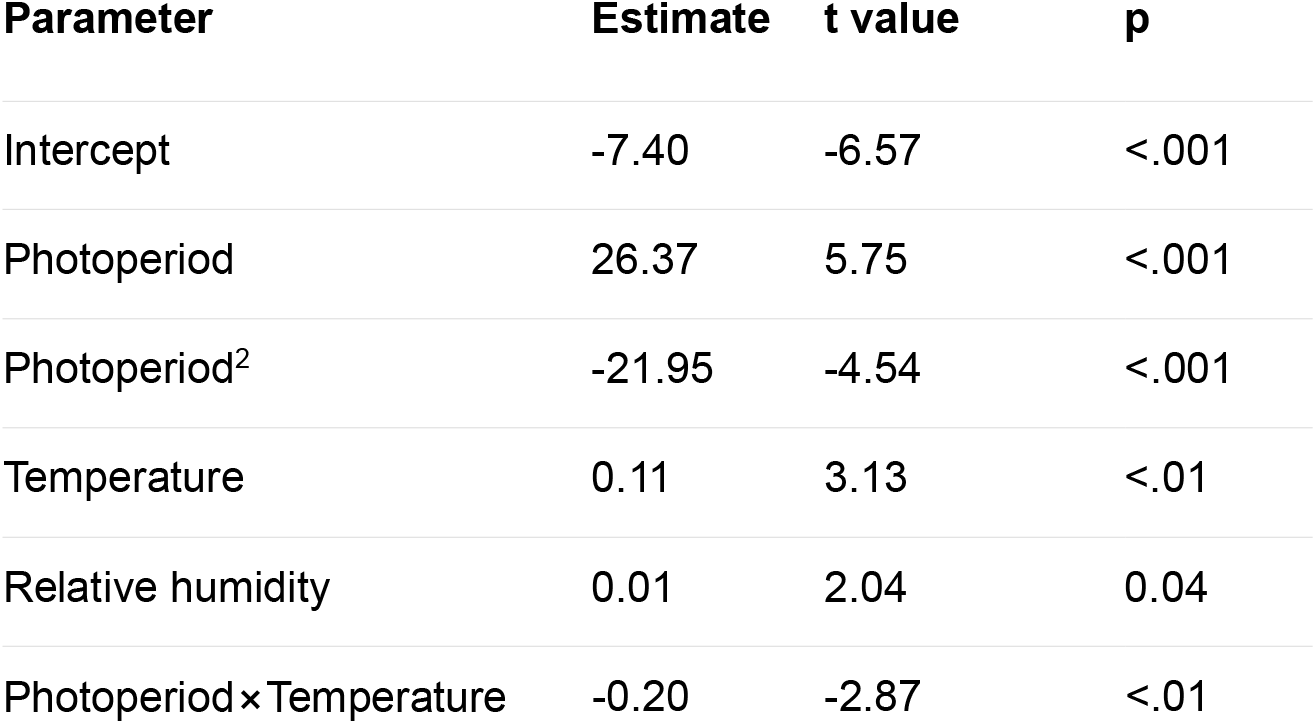
Parameter values and significance for Model 4.

| Parameter | Estimate | t value | p |
| --- | --- | --- | --- |
| Intercept | -7.40 | -6.57 | <.001 |
| Photoperiod | 26.37 | 5.75 | <.001 |
| Photoperiod<sup>2</sup> | -21.95 | -4.54 | <.001 |
| Temperature | 0.11 | 3.13 | <.01 |
| Relative humidity | 0.01 | 2.04 | 0.04 |
| Photoperiod × Temperature | -0.20 | -2.87 | <.01 |

## Discussion

We used passive acoustic monitoring to continuously record calling activity in *B. pulchella* inhabiting a permanent pond in a subtropical region of South America, while simultaneously recording ambient environmental cues. Chorusing activity followed a seasonal pattern, with more prolonged activity during warmer months. Calling activity correlated distinctly with temperature and photoperiod, exhibiting a significant interaction between these variables, whereas no significant association was found with rainfall. In addition, our research validated semi-automated data processing techniques for PAM, contributing to the limited number of such methodologies available for studying anuran choruses in southern South America.

With our use of passive recording systems and continuous acoustic monitoring, we were able to precisely characterize daily vocalization patterns. We found that individuals modulate the onset of calling activity in response to seasonal changes and adjust the timing of chorus initiation and termination according to time of day, season, temperature, and photoperiod. Similar patterns have been reported in other animal taxa (Åsvestad et al., 2024; Hao et al., 2025; Singer et al., 2025). The peak hour of chorus activity in *B. pulchella* does not appear to correlate with photoperiod, but this association varies among frog species. Some anuran species adjust vocal peaks seasonally—linked to water availability, rainfall, or predation risk—while others show no variation throughout the year (Bertoluci and Rodrigues, 2002; Gottsberger and Gruber, 2004). These species differences are likely due to the geographic location of the study populations and to variation in active recording methods, which are often sporadic and less precise.

Temporal adjustments in calling activity may serve multiple adaptive functions. Modifying calling schedules could minimize acoustic interference with other anuran species or sympatric organisms, thereby optimizing signal transmission within the soundscape (Brooke et al., 2000). Additionally, it may depend on the presence of conspecifics, including the time it takes for them to arrive or begin calling. Furthermore, these shifts may function as an antipredator strategy, reducing exposure to predators that rely on acoustic cues for prey detection. Beyond these selective pressures, temporal variation in vocal behavior may also influence reproductive success, as female arrival times fluctuate seasonally (Ryan et al., 1981). These seasonal dynamics may further align with environmental constraints, such as the need for optimal water temperatures to support successful egg deposition and embryonic development (Brinkløv et al., 2023).

The influence of environmental cues on acoustic signal emission has been extensively studied for decades, however much of this research has relied on sporadic active recordings. More recently, PAM has emerged as a valuable tool to investigate population-level vocal behavior in vertebrates—such as in bats (Brinkløv et al., 2023) and birds (Rumelt et al., 2021)—offering advantages in cost-efficiency, autonomy, and temporal precision. This approach enables extended monitoring periods and the simultaneous study of multiple species. In anurans, PAM has revealed high acoustic diversity and species-specific calling patterns, particularly in long-term studies conducted in tropical and subtropical regions (Cañas et al., 2023; Sugai et al., 2019). However, studies in temperate regions of southern South America remain scarce, despite the potential of these systems to yield novel insights into vocal behavior dynamics (Sugai et al., 2019). Here, we investigated a species that exhibits predominant calling activity year-round in a permanent pond.

To date, few studies have quantified calling activity at minute-scale temporal resolution over a full annual cycle while concurrently recording environmental variables in a subtropical temperate zone (Boullhesen et al., 2019). While this approach facilitates detailed and continuous data collection, it also poses significant challenges due to the volume and complexity of the resulting datasets. Automated call detection using signal-to-noise ratio thresholds, validated through manual inspection, enables reliable identification of calling presence. However, this method provides only coarse estimates of call abundance, limiting its utility for finer-scale quantification (Metcalf et al., 2024).

Understanding how environmental cues—such as temperature, humidity, rainfall, and photoperiod—shape natural anuran populations provide valuable insight into their reproductive activity and, over time, their life history. Traditionally, rainfall and humidity have been considered primary factors influencing calling activity in the tropics, whereas temperature and photoperiod show stronger correlations with vocal behavior than in temperate regions (Duellman & Trueb, 1994). In temperate climates, this may stem from the fact that rainfall occurs year-round and is not a distinct seasonal variable. However, literature suggests that these environmental cues—particularly rainfall and temperature—are not always directly linked to reproductive behavior or calling activity in specific tropical and subtropical anuran populations (Akmentins et al., 2023, 2015; Ospina et al., 2013).

This diversity has been documented among species of the genus *Boana*. For example, a tropical Boana population in Brazil showed no association between vocal activity and temperature (Bonnefond et al., 2020), whereas Canavero et al. (2008) reported a positive association in a subtropical temperate population in Uruguay. Regarding rainfall, increased vocal activity during the rainy season compared to the dry season has been observed in tropical populations of the genus Boana in Brazil, suggesting a seasonal pattern in call emission. However, these studies did not assess whether this pattern is directly correlated with rainfall or temperature (Duarte et al., 2019). Conversely, a positive correlation between vocal activity and rainfall has been reported in a subtropical population in Uruguay (Da Rosa, 2020). In contrast, no such association was found in populations from southeastern Brazil and Uruguay (Both et al., 2008; Canavero et al., 2008).

Unlike previous research, we found no correlation between calling activity and rainfall, which further supports the lack of an effect of atmospheric pressure—a well-known predictor of precipitation (Oseen and Wassersug, 2002; Plenderleith et al., 2018). Several factors may account for this discrepancy. First, previous studies relied on sporadic active monitoring and collected data during a short time window each day, resulting in a limited dataset. Second, the breeding ponds were not permanent features. Finally, the geographic location of the study area in Rosa (2020) differs from that of the present research. These contextual and methodological differences likely explain the contrasting results, highlighting the importance of standardized approaches and site-specific evaluations in acoustic monitoring studies.

The effect of photoperiod on calling behavior is not well studied, but it has been identified as a key determinant of seasonal anuran activity, particularly in temperate regions (Both et al., 2008; Canavero et al., 2008; Canavero and Arim, 2009). Our findings are consistent with those of Canavero and Arim (2009), who applied a structural equation model to demonstrate a strong correlation between photoperiod and anuran species richness in calling activity. Unlike our study, which focused on a single species with continuous monitoring, Canavero et al. (2008) analyzed multiple species concurrently and relied on active, sporadic monitoring.

A relevant finding from our study indicates that temperature predicts calling behavior differently depending on photoperiod and average temperature (Figure 3). This highlights plasticity in the adjustment of calling activity in response to day length, suggesting a seasonal modulation that expands or restricts the thermal range in which calling occurs. This result aligns with expectations for species inhabiting temperate zones, where moderate annual variation in the thermal regime is typical (Llusia et al., 2013). This finding not only reflects the well-established thermal responsiveness of ectotherms, but also provides evidence of their capacity to modulate vocal behavior in response to photoperiodic variation (Oseen and Wassersug, 2002).

Collectively, these findings underscore the complex interactions between environmental drivers, reproductive timing, and acoustic communication in tropical anurans. Because environmental cues may exert divergent effects depending on local conditions and study designs, high temporal resolution is essential for capturing the fine-scale dynamics of vocal behavior. Passive acoustic monitoring offers a powerful tool for exploring detailed temporal dynamics across climatic gradients. The consistent role of thermal and photic cycles as key regulators of calling behavior points to underlying physiological and adaptive mechanisms that merit further investigation. Detailed characterization of species-specific vocal patterns will be essential to better understand the ecological significance and evolutionary trajectories of these vocal adjustments under changing environmental conditions.

## Acknowledgements

We sincerely thank the field owners, Álvaro Mazzilli Millán and his family, as well as Mercedes Rodríguez and Isidro Fernández, for their generous hospitality. Ana Silva, Carlos Colacce and La Comarca provided housing. We also extend our gratitude to Esteban Russi for his invaluable assistance in the field and to Mikaela Cúparo for her dedication in manually validating the data. We thank Kent Dunlap and Kim Hoke for their helpful reading and constructive feedback on this article.

## References

Akmentins, M.S., Boullhesen, M., Vaira, M., Pereyra, L.C., 2023. Vocalization Behavior and Calling Phenology in Two Direct-developing Frogs of the genus Oreobates of Yungas Andean Forests in Northwest Argentina.

Akmentins, M.S., Pereyra, L.C., Sanabria, E.A., Vaira, M., 2015. Patterns of daily and seasonal calling activity of a direct-developing frog of the subtropical Andean forests of Argentina. Bioacoustics 24, 89–99. 10.1080/09524622.2014.965217

Åsvestad, L., Ahonen, H., Menze, S., Lowther, A., Lindstrøm, U., Krafft, B.A., 2024. Seasonal acoustic presence of marine mammals at the South Orkney Islands, Scotia Sea. Royal Society Open Science 11, 230233. 10.1098/rsos.230233

Barkan, C.L., Leininger, E.C., Zornik, E., 2021. Everything in Modulation: Neuromodulators as Keys to Understanding Communication Dynamics. Integrative and Comparative Biology 61, 854–866. 10.1093/icb/icab102

Basso, N.G., Basso, G., 1987. Análisis acústico del canto nupcial de Hyla pulchella pulchella Dumeril & Bibron 1841 (Anura: Hylidae). An Mus Hist Nat Valparaiso 18, 109–114.

Bertoluci, J., Rodrigues, M.T., 2002. Seasonal patterns of breeding activity of Atlantic Rainforest anurans at Boracéia, Southeastern Brazil. 10.1163/156853802760061804

Bonnefond, A., Courtois, E.A., Sueur, J., Sugai, L.S.M., Llusia, D., 2020. Climatic breadth of calling behaviour in two widespread Neotropical frogs: Insights from humidity extremes. Global Change Biology 26, 5431–5446. 10.1111/gcb.15266

Both, C., Kaefer, í.L., Santos, T.G., Cechin, S.T.Z., 2008. An austral anuran assemblage in the Neotropics: seasonal occurrence correlated with photoperiod. Journal of Natural History 42, 205–222. 10.1080/00222930701847923

Boullhesen, M., Salica, M.J., Pereyra, L.C., Akmentins, M.S., 2019. Actividad vocal diaria y su relación con claves ambientales en un ensamble de anuros en las Yungas de Jujuy, Argentina. Cuad. Herpetol. 33. 10.31017/CdH.2019.(2019-012)

Brenowitz, E.A., Rose, G., Capranica, R.R., 1985. Neural correlates of temperature coupling in the vocal communication system of the gray treefrog (Hyla versicolor). Brain Res 359, 364–367. 10.1016/0006-8993(85)91452-0

Brinkløv, S.M.M., Macaulay, J., Bergler, C., Tougaard, J., Beedholm, K., Elmeros, M., Madsen, P.T., 2023. Open-source workflow approaches to passive acoustic monitoring of bats. Methods in Ecology and Evolution 14, 1747–1763. 10.1111/2041-210X.14131

Brodie, S., Allen-Ankins, S., Schwarzkopf, L., 2025. Environmental influences on chorusing patterns in an Australian tropical savanna frog community. Ecosphere 16, e70153. 10.1002/ecs2.70153

Brooke, P.N., Alford, R.A., Schwarzkopf, L., 2000. Environmental and social factors influence chorusing behaviour in a tropical frog: examining various temporal and spatial scales. Behav Ecol Sociobiol 49, 79–87. 10.1007/s002650000256

Cañas, J.S., Toro-Gómez, M.P., Sugai, L.S.M., Benítez Restrepo, H.D., Rudas, J., Posso Bautista, B., Toledo, L.F., Dena, S., Domingos, A.H.R., De Souza, F.L., Neckel-Oliveira, S., Da Rosa, A., Carvalho-Rocha, V., Bernardy, J.V., Sugai, J.L.M.M., Dos Santos, C.E., Bastos, R.P., Llusia, D., Ulloa, J.S., 2023. A dataset for benchmarking Neotropical anuran calls identification in passive acoustic monitoring. Sci Data 10, 771. 10.1038/s41597-023-02666-2

Canavero, A., Arim, M., 2009. Clues supporting photoperiod as the main determinant of seasonal variation in amphibian activity. Journal of Natural History 43, 2975–2984. 10.1080/00222930903377539

Canavero, A., Arim, M., Naya, D.E., 2008. Calling activity patterns in an anuran assemblage : the role of seasonal trends and weather determinants. North-Western Journal of Zoology 4, 29–41.

Chmura, H.E., Kharouba, H.M., Ashander, J., Ehlman, S.M., Rivest, E.B., Yang, L.H., 2019. The mechanisms of phenology: the patterns and processes of phenological shifts. Ecological Monographs 89, e01337. 10.1002/ecm.1337

Da Rosa, I., 2020. Environmental Determinants of Calling Activity and Temporal Body Size Variation in Males of Boana pulchella (Anura: Hylidae) from Southern Uruguay. South American Journal of Herpetology 2020, 1. 10.2994/SAJH-D-17-00033.1

Duarte, M.H.L., Caliari, E.P., Viana, Y.P., Nascimento, L.B., 2019. A natural orchestra: how are anuran choruses formed in artificial ponds in southeast Brazil? Amphib.-Reptilia 40, 373–382. 10.1163/15685381-20191079

Duellman, W.E., Trueb, L., 1994. Biology of Amphibians. JHU Press.

Feng, A.S., Riede, T., Arch, V.S., Yu, Z., Xu, Z., Yu, X., Shen, J., 2009. Diversity of the Vocal Signals of Concave-Eared Torrent Frogs (Odorrana tormota): Evidence for Individual Signatures. Ethology 115, 1015–1028. 10.1111/j.1439-0310.2009.01692.x

Gentner, T.Q., Margoliash, D., 2003. The Neuroethology of Vocal Communication: Perception and Cognition, in: Simmons, A.M., Fay, R.R., Popper, A.N. (Eds.), Acoustic Communication, Springer Handbook of Auditory Research. Springer-Verlag, New York, pp. 324–386. 10.1007/0-387-22762-8_7

Gerhardt, H.C., 2001. Acoustic communication in two groups of closely related treefrogs, in: Advances in the Study of Behavior. Academic Press, pp. 99–167. 10.1016/S0065-3454(01)80006-1

Gibb, R., Browning, E., Glover-Kapfer, P., Jones, K.E., 2019. Emerging opportunities and challenges for passive acoustics in ecological assessment and monitoring. Methods Ecol Evol 10, 169–185. 10.1111/2041-210x.13101

Gottsberger, B., Gruber, E., 2004. Temporal partitioning of reproductive activity in a neotropical anuran community. Journal of Tropical Ecology 20, 271–280. 10.1017/S0266467403001172

Hao, P., Rao, X., Liang, W., Zhang, Y., 2025. Temporal patterns and environmental drivers of the red junglefowl vocalization. BMC Ecol Evo 25, 54. 10.1186/s12862-025-02391-x

Hauselberger, K.F., Alford, R.A., 2005. EFFECTS OF SEASON AND WEATHER ON CALLING IN THE AUSTRALIAN MICROHYLID FROGS AUSTROCHAPERINA ROBUSTA AND COPHIXALUS ORNATUS. Herpetologica 61, 349–363. 10.1655/04-03.1

Hawkins, L.A., Parsons, M.J.G., McCauley, R.D., Parnum, I.M., Erbe, C., 2025. Passive acoustic monitoring of fish choruses: a review to inform the development of a monitoring and management tool. Rev Fish Biol Fisheries 35, 847–874. 10.1007/s11160-025-09936-9

Lin, F.-C., Lin, S.-M., Godfrey, S.S., 2024. Hidden social complexity behind vocal and acoustic communication in non-avian reptiles. Philosophical Transactions of the Royal Society B: Biological Sciences 379, 20230200. 10.1098/rstb.2023.0200

Llusia, D., Márquez, R., Beltrán, J.F., Benítez, M., do Amaral, J.P., 2013. Calling behaviour under climate change: geographical and seasonal variation of calling temperatures in ectotherms. Global Change Biology 19, 2655–2674. 10.1111/gcb.12267

Maneyro, R., Carreira, S., 2012. Guía de Anfibios del Uruguay. Ediciones de la Fuga, Montevideo.

Metcalf, O.C., Nunes, C.A., Abrahams, C., Baccaro, F.B., Bradfer-Lawrence, T., Lees, A.C., Vale, E.M., Barlow, J., 2024. The efficacy of acoustic indices for monitoring abundance and diversity in soil soundscapes. Ecological Indicators 169, 112954. 10.1016/j.ecolind.2024.112954

Narins, P., Gerhardt, H.C., Christensen-Dalsgaard, J., 2023. A Nasty, Brutish, and Short History of Amphibian Bioacoustics. pp. 75–112. 10.1007/978-3-031-41320-9_4

Narins, P.M., Meenderink, S.W.F., 2014a. Climate change and frog calls: long-term correlations along a tropical altitudinal gradient. Proceedings of the Royal Society B: Biological Sciences 281, 20140401. 10.1098/rspb.2014.0401

Narins, P.M., Meenderink, S.W.F., 2014b. Climate change and frog calls: long-term correlations along a tropical altitudinal gradient. Proc. R. Soc. B. 281, 20140401. 10.1098/rspb.2014.0401

Navas, C.A., 1996. The effect of temperature on the vocal activity of tropical anurans: a comparison of high and low-elevation species. Journal of Herpetology 488–497.

Oestreich, W.K., Oliver, R.Y., Chapman, M.S., Go, M.C., McKenna, M.F., 2024. Listening to animal behavior to understand changing ecosystems. Trends in Ecology & Evolution 39, 961–973. 10.1016/j.tree.2024.06.007

Oseen, K.L., Wassersug, R.J., 2002. Environmental factors influencing calling in sympatric anurans. Oecologia 133, 616–625. 10.1007/s00442-002-1067-5

Ospina, O.E., Villanueva-Rivera, L.J., Corrada-Bravo, C.J., Aide, T.M., 2013. Variable response of anuran calling activity to daily precipitation and temperature: implications for climate change. Ecosphere 4, art47. 10.1890/ES12-00258.1

Pauly, G.B., Bernal, X.E., Rand, A.S., Ryan, M.J., 2006. The Vocal Sac Increases Call Rate in the Túngara FrogPhysalaemus pustulosus. Physiological and Biochemical Zoology 79, 708–719. 10.1086/504613

Plenderleith, T.L., Stratford, D., Lollback, G.W., Chapple, D.G., Reina, R.D., Hero, J.-M., 2018. Calling phenology of a diverse amphibian assemblage in response to meteorological conditions. Int J Biometeorol 62, 873–882. 10.1007/s00484-017-1490-2

Posit team, 2025. RStudio: Integrated Development Environment for R. Posit Software, PBC, Boston, MA.

R Core Team, 2025. R: A Language and Environment for Statistical Computing. R Foundation for Statistical Computing, Vienna, Austria.

Ribeiro, J.W., Sugai, L.S.M., Campos-Cerqueira, M., 2017. Passive acoustic monitoring as a complementary strategy to assess biodiversity in the Brazilian Amazonia. Biodivers Conserv 26, 2999–3002. 10.1007/s10531-017-1390-0

Rodriguez-Santiago, M., Russi, Esteban, Zornik, E., Pouso, Paula, Hoke, K.L., 2023. Variability in the vocal repertoire of a South American treefrog. Presented at the Society for Integrative Comparative Biology, Annual Meeting, United States.

Rumelt, R.B., Basto, A., Mere Roncal, C., 2021. Automated audio recording as a means of surveying tinamous (Tinamidae) in the Peruvian Amazon. Ecol Evol 11, 13518–13531. 10.1002/ece3.8078

Ryan, M.J., Tuttle, M.D., Taft, L.K., 1981. The Costs and Benefits of Frog Chorusing Behavior. Behavioral Ecology and Sociobiology 8, 273–278.

Schalk, C.M., Saenz, D., 2016. Environmental drivers of anuran calling phenology in a seasonal Neotropical ecosystem. Austral Ecology 41, 16–27. 10.1111/aec.12281

Simmons, A., Fay, R.R., 2006. Acoustic Communication. Springer Science & Business Media.

Singer, D., Kamp, J., Hondong, H., Schuldt, A., Hagge, J., 2025. Diel and seasonal vocal activity patterns revealed by passive acoustic monitoring suggest expert recommendations for breeding bird surveys need adjustment. J Ornithol 1–16. 10.1007/s10336-025-02307-y

Sueur, J., Aubin, T., Simonis, C., 2008. Seewave, a Free Modular Tool for Sound Analysis and Synthesis. Bioacoustics 18, 213–226. 10.1080/09524622.2008.9753600

Sugai, L.S.M., Silva, T.S.F., Llusia, D., Siqueira, T., 2021. Drivers of assemblage-wide calling activity in tropical anurans and the role of temporal resolution. Journal of Animal Ecology 90, 673–684. 10.1111/1365-2656.13399

Sugai, L.S.M., Silva, T.S.F., Ribeiro, J.W., Jr, Llusia, D., 2019. Terrestrial Passive Acoustic Monitoring: Review and Perspectives. BioScience 69, 15–25. 10.1093/biosci/biy147

Visser, M.E., Caro, S.P., van Oers, K., Schaper, S.V., Helm, B., 2010. Phenology, seasonal timing and circannual rhythms: towards a unified framework. Philosophical Transactions of the Royal Society B: Biological Sciences 365, 3113–3127. 10.1098/rstb.2010.0111

Young, B., Mathevon, N., Tang, Y., 2013. Reptile Auditory Neuroethology: What Do Reptiles Do with Their Hearing? pp. 323–346. 10.1007/2506_2013_30

Ziegler, L., Arim, M., Bozinovic, F., 2016. Intraspecific scaling in frog calls: the interplay of temperature, body size and metabolic condition. Oecologia 181, 673–681. 10.1007/s00442-015-3499-8

Ziegler, L., Arim, M., Narins, P.M., 2011. Linking amphibian call structure to the environment: The interplay between phenotypic flexibility and individual attributes. Behavioral Ecology 22, 520–526. 10.1093/beheco/arr011

