## Supplementary material for "Environmental Drivers of Calling Activity in a Southern Subtropical Anuran Assemblage: Insights from Passive Acoustic Monitoring": Figures S1 and S2

Supplementary Figure S1- Pouso et al

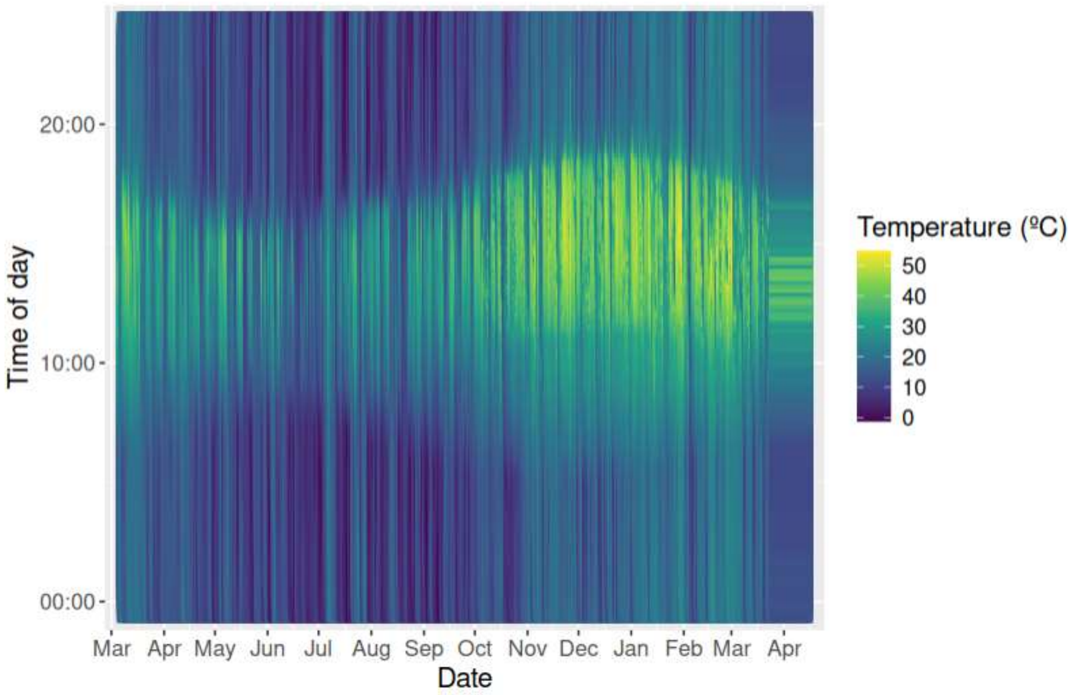


Supplementary Figure S1. Seasonal and hourly variation in temperature over a year. Warmer conditions (yellow regions) are concentrated during daytime hours, while cooler temperatures (blue shades) prevail at night. (I don't see anything about light intensity here. Also, change months to English or to numbers.)

Supplementary Figure S2-Pouso et al

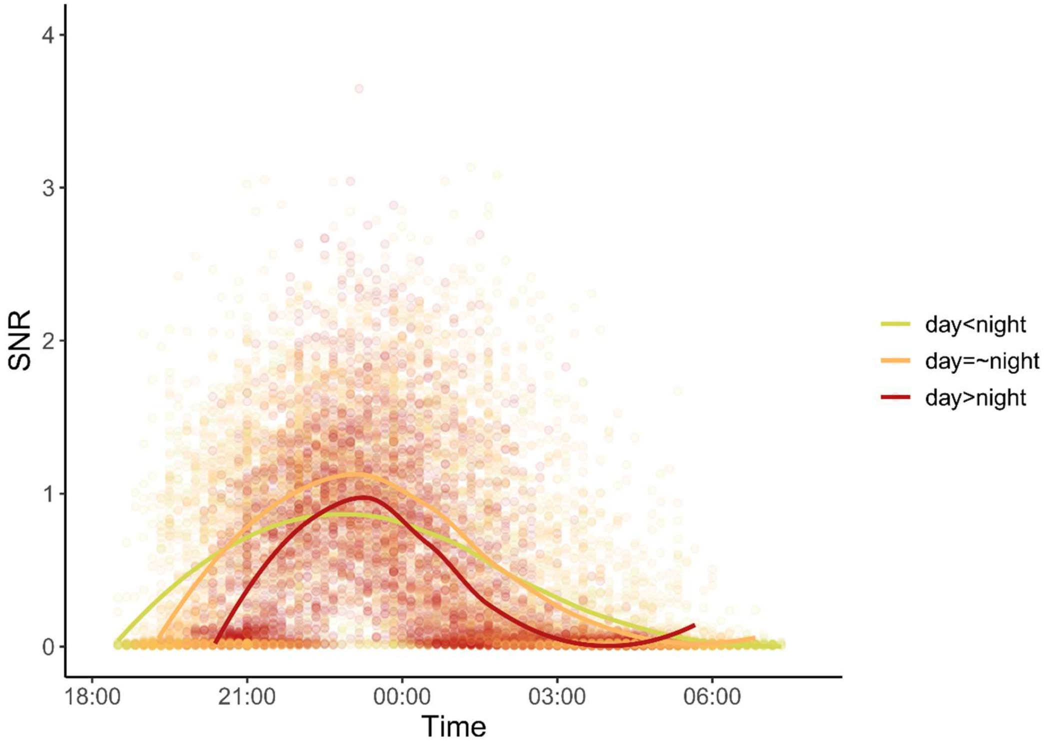


Supplementary Figure S2. Calling activity profiles by time and photoperiod. Time at peak values for each season are (median + IQR): day<night: 11:30 PM (240 mins); day=~night: 10:45 PM (108 mins); day>night: 11:00 PM (70 mins). Time at peak medians did not differ significantly between photoperiods (p-values > .4).
